# Scale dependent patterns in interaction diversity maintain resiliency in a frequently disturbed ecosystem

**DOI:** 10.1101/528745

**Authors:** Jane E. Dell, Danielle M. Salcido, Will Lumpkin, Lora A. Richards, Scott M. Pokswinski, E. Louise Loudermilk, Joseph J. O’Brien, Lee A. Dyer

**Affiliations:** Ecology, Evolution, and Conservation Biology, Department of Biology, University of Nevada, Reno, Reno, NV USA; Wildland Fire Science Program, Tall Timbers Research Station, Tallahassee, FL USA; Center for Forest Disturbance Science, Southern Research Center, US Forest Service, Athens, GA, USA

**Keywords:** interaction diversity, scale-dependency, response diversity, resilience, tri-trophic interaction, *Pinus palustris*, prescribed fire

## Abstract

Frequently disturbed ecosystems are characterized by resilience to ecological disturbances. For example, longleaf pine ecosystems are exposed to frequent fire disturbance, and this feature sustains biodiversity. We examined how fire frequency maintains beta diversity of multi-trophic interactions, as this community parameter provides a measure of functional redundancy of an ecosystem. We found that turnover in interaction diversity at small local scales is highest in the most frequently burned stands, conferring immediate resiliency to disturbance by fire. Interactions become more specialized and less resilient as fire frequency decreases. Local scale patterns of interaction diversity contribute to broader scale patterns and confer long-term ecosystem resiliency. Such natural disturbances are likely to be important for maintaining regional diversity of interactions for a broad range of ecosystems.

## Introduction

Disturbances are significant features of ecosystems, with frequency and intensity being important for shaping not only community composition, structure, and function but also serving as selective forces in the evolution of life history strategies, especially in disturbance-prone ecosystems (Sousa 1984, Seidl et al. 2016). Resiliency and high beta diversity are critical features of many of these frequently disturbed ecosystems (Elmqvist et al. 2003, Larson et al. 2013). Regular disturbance events maintain a diverse and functional ecosystem state in disturbance-dependent systems and within this context, resiliency is defined as ecosystem recovery to pre-disturbance levels of parameters such as diversity, population measures, and nutrient cycling immediately post-disturbance. Conversely, a disruption of the disturbance regime, such as reduced frequency, represents a transformational and longer-term perturbation where ecosystem structure and function shifts and pushes such systems to unpredictable or unstable, alternative states (Beisner et al. 2003, Bowman et al. 2016). In this context, removal of disturbance erodes long-term resilience of a disturbance-adapted ecosystem.

Resiliency requires a minimum level of underlying species and functional diversities to allow for multiple pathways towards post-disturbance responses (Seidl et al. 2016). The return to pre-disturbance function due to functional redundancies provided by biological diversity is also known as response diversity (Figure 1). Elmqvist et al. (2003) define response diversity as the diversity of responses to disturbance among different assemblages of species that contribute to equivalent ecosystem functions. However, response diversity is not simply equivalent to species richness for different broad taxa or at different trophic levels, because ecological communities are comprised of species that interact in different functional ways. For instance, a broad diet breadth or shared basal resources provide functional redundancy and are indicative of response diversity and trophic network stability (Pilosof et al. 2017; Figure 1). As such, quantification of response diversity requires easily measured metrics, such as interaction diversity, in order to understand the effects of disturbance on the interactions between species (Dyer et al. 2010). Critical ecosystem functions, such as pollination, population control of herbivores by natural enemies, and seed dispersal are dependent upon a broad range of biotic interactions at small scales, the loss of which can precipitate species extinctions and loss of ecological function (Kremen et al. 2007, Valiente-Banuet et al. 2015; Figure 1). Therefore, it is also important to consider interaction diversity, defined as the richness and relative abundance of species interactions in a community (Dyer et al. 2010), as a primary contributor to ecosystem resilience and a critical component of response diversity. While species richness and potential interactions are necessarily positively correlated, diversity of species and diversity of interactions can have different effects on ecosystem function and stability (Pardikes et al. 2018). Like other diversity metrics, interaction diversity across the landscape has alpha, beta, and gamma components that can differ substantially from species diversity.

**Figure 1:**
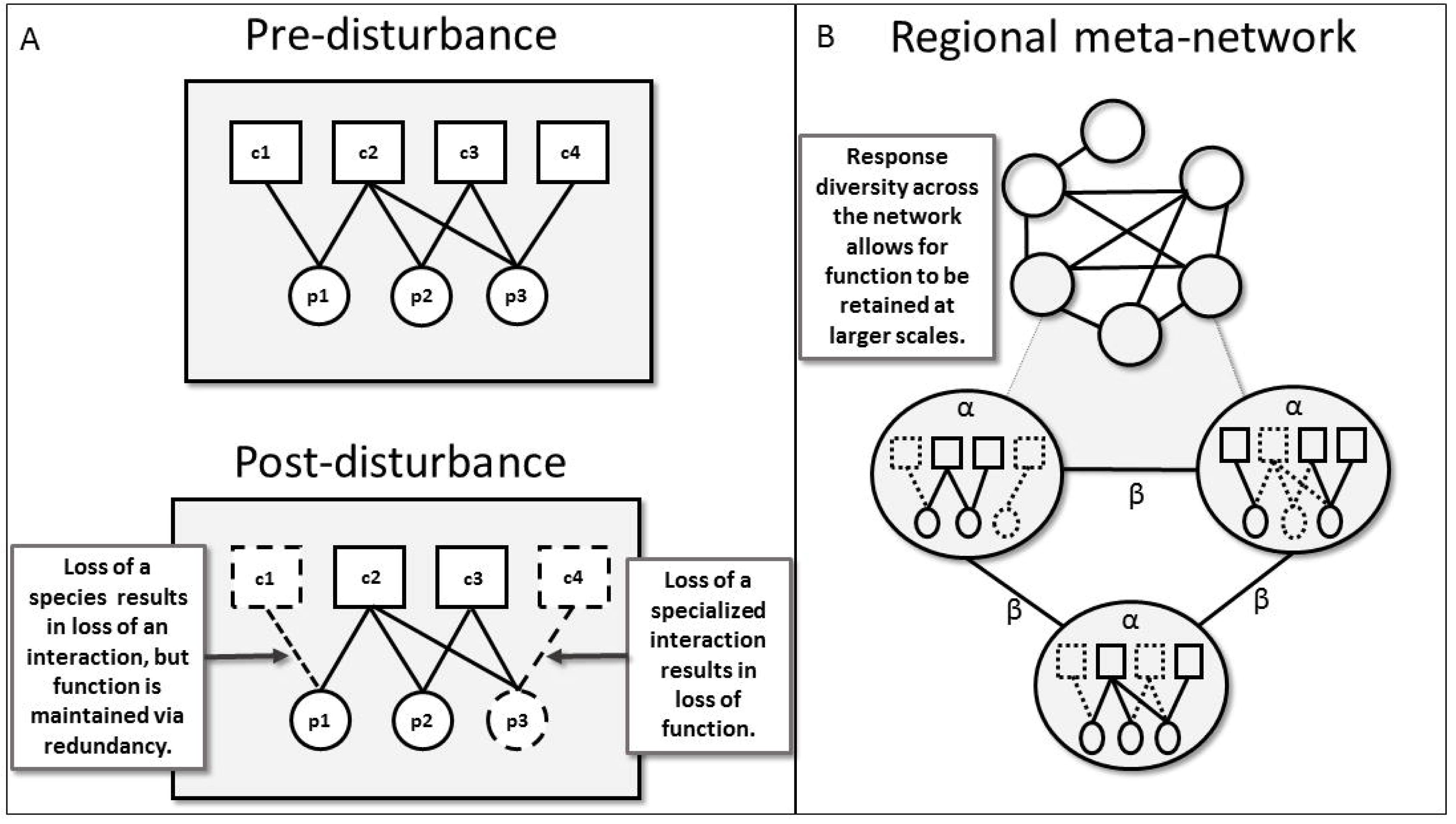
**(A)** Pre-disturbance and post-disturbance metawebs, displaying the full regional pool of species and potential interactions. Here nodes represent individual species of primary producers (circles) and herbivores (squares), while edges (links) represent interactions between species. Post-disturbance, the loss of species and interactions are indicated by dashed edges. In this case the loss of a single species (c1) also results in the loss of an interaction, however ecological function (i.e. nutrient cycling via consumption of this plant species) is maintained as a redundant interaction occurs with another herbivore species (c2). Conversely, the loss of a specialized interaction may result in the loss of ecological function. In this example, the interaction between a plant (p3) and herbivore (c4) no longer occurs, reducing functional diversity and eventual loss of partner species (p3-c4). **(B)** The regional meta-network, for which nodes represent plots and edges represent shared interactions between plots. Three plots are enlarged so that we may examine the corresponding local networks of interactions. While α-diversity of species and interactions are calculated within each plot, β-diversity is calculated between plots. Focusing in on shared interactions between three individual plots illustrates the turnover of interactions between local plots (high β-diversity), this β-diversity summarizes variation in post-disturbance responses, which provides ecological resiliency.

As with any metric exploring patterns of biodiversity, concepts of scale are necessary to consider when examining the causes and consequences of the richness and turnover of interacting species (Giron et al. 2018). Patterns observed at larger scales represent the combined processes occurring at smaller scales, but it is not always clear how patterns at nested scales relate to one another. Species richness differs among local and regional scales (Witman et al. 2004, Rahbek and Graves 2001) partly because regional and local diversity are shaped by different processes. For plant-insect networks, regional processes are affected more by large-scale evolutionary and historical factors, such as speciation, dispersal, extinction, and biogeographical history, while local processes include ecological effects such as, biotic interactions, resource availability, and disturbance. Furthermore, interaction networks are not static, and patterns in interaction diversity are unlikely to be constant across the landscape or at different spatial extents. At smaller local scales, trait-distributions, environmental conditions and species abundance will affect the potential of two co-occurring species to interact (Poisot 2015). Regionally, interaction diversity values can change substantially depending on the scale at which they are examined (Pardikes et al. 2018).

In this study we focused on trophic interactions between host plants, arthropod herbivores, and their parasitoid enemies in a frequently disturbed ecosystem across a large fire-adapted forest ecosystem. Disturbance by fire has been a part of terrestrial ecosystems since the Silurian Period and is an essential process for maintaining both ecosystem function and biological diversity in fire dependent ecosystems (Pausas and Keeley 2009), such as the frequently burned longleaf pine (*Pinus palustris* Mill., Kirkman et al. 2004, O’Brien et al. 2008, Mitchell et al. 2009). In the absence of fire, competitive advantage is given to faster growing, non-fire dependent broadleaved vegetation, resulting in a closed canopy, extensive habitat degradation, and reductions in plant diversity (Mitchell et al. 2009, Noss et al. 2014). The removal of fire from the landscape initiates a shifting ecosystem trajectory where fire-adapted species are replaced by other species assemblages, yielding an alternative stable state (Beisner et al. 2003).

Our primary objective was to quantify interaction diversity across a time since fire gradient, in order to assess the effect of longer fire return intervals on biotic community interactions and potential for resiliency in longleaf pine forests. We posit that resiliency will be greatest in ecosystems where there is functional redundancy, (i.e. high response diversity), and that this functional redundancy is greatest when levels of beta interaction diversity (for multiple scales) are maintained (Figure 1). Higher levels of turnover in interactions are indicative of increased ecological function (Lepesqueur et al. 2018) such that a reduction in beta diversity represents a homogenization of interactions which may reduce ecosystem function by affecting productivity, resilience to disturbance, and vulnerability to biological invasion (Balata et al. 2007, Dell et al. 2019). As frequent fire maintains high-levels of plant diversity and ecosystem function, we predict that large-scale interaction diversity will be higher in frequently burned stands than in stands with longer times since fire. Second, to understand the way interaction diversity varies with scale, we investigated how these patterns vary at both the small, plot-level versus broader, regional-level scales. Many understory plant species have a patchy distribution in longleaf pine because of fine-scale variation in fuel and fire heterogeneity (Menges and Hawkes 1998, Dell et al. 2017), and diversity of these plants is best quantified at small spatial scales, therefore we expect that interaction diversity will also vary and patterns will change with increasing spatial scale. Due to the connectivity between these spatial scales any such local scale patterns of interaction diversity will contribute to broader scale patterns and confer long-term ecosystem resiliency for the region.

## Materials and methods

### Study area

Research was conducted in longleaf pine forests across the Gulf Coastal Plain during 2013 to 2016. Sites included Eglin Air Force Base and Blackwater River State Forest located in the Florida panhandle and Solon Dixon Forestry Education Center and Conecuh National Forest in southern Alabama. The fire regime in longleaf pine ecosystems is characterized by high-frequency, low-intensity surface fires with return intervals of 1-5 years (Mitchell et al. 2009). Numerous longleaf pine stands within the region are actively managed by prescribed fires with a target of an 18-month to two-year fire return interval (Hiers et al. 2007). However, there exist stands within all management areas that have not experienced burning for longer periods of time including up to several decades. Sampling includes both frequently burned and infrequently burned areas as well as an intermediate transitional state.

### Field collection

We established sixty-seven, 30-m diameter plots in forested stands that varied in the time since last disturbance by fire (Figure 2). Based on available fire history records and vegetative indicator species associated with known fire return intervals, plots were placed into a burn category; frequently burned (fire return interval (FRI): 1-5 years, n = 49), intermediately burned (FRI: 5-25 years, n = 9), and infrequently burned (FRI: >25 years, n = 9).

**Figure 2:**
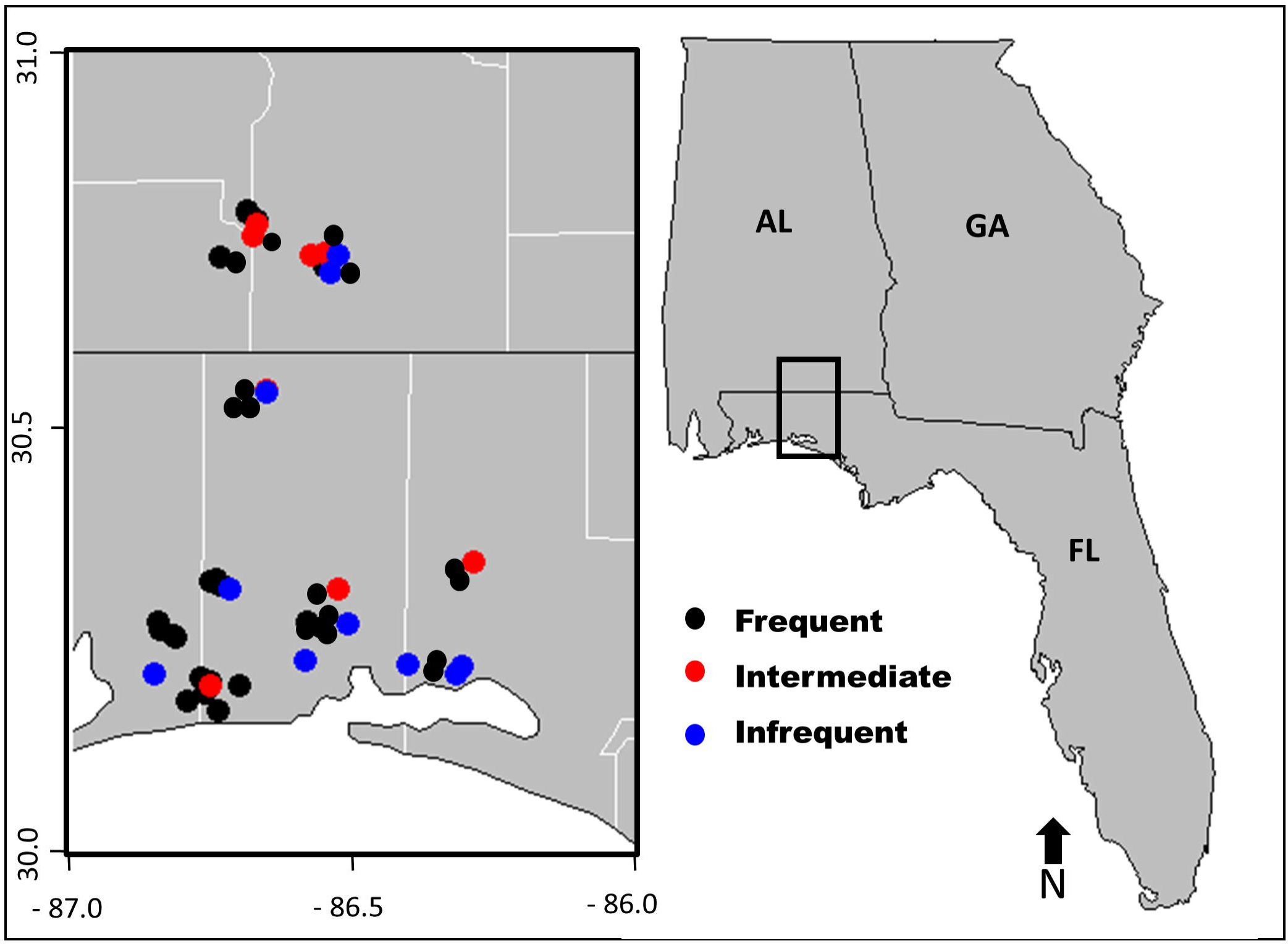
Reference map of Southeastern United States with study area highlighted in black rectangle. Within the study area, individual plot locations are indicated by colored dots corresponding to each of the three fire return intervals.

Throughout the entire growing season, Lepidopteran larvae were sampled within each plot using beat sheets and visual searches in a standardized format moving clockwise around the plot. Within each fire return interval type, we also generally collected caterpillars outside of standardized plots to further expand the trophic network within each fire return interval category. Caterpillars were reared out to adulthood or eclosion of a parasitoid. Host plant associations were based on the vegetation from which the caterpillars were collected and confirmed through feeding in the laboratory. Host plants and arthropods were identified to species or were assigned a morphotype based on morphological characteristics, behavior, and host plant record following Wagner (2005). Sampled arthropods were deposited into the research collection at the University of Nevada, Reno Museum of Natural History.

### Quantification of diversity

Diversity was estimated for species and interactions at two scales; the plot-level or local scale and the broader, regional-level scale. The regional level is defined as the aggregate of all plots and general collecting sites within each fire return interval category over the entire range of the study. It should be noted that longleaf pine forests in this study may have frequently burned stands adjacent to infrequently burned stands, therefore our use of the term regional does not infer a singular spatial aggregation, but rather a broader character-state organization. Interaction diversity was based on the richness and abundance of interactions between species, where richness is the number of unique interactions and abundance the total number of each interaction (Dyer et al. 2010; Figure 3). Bipartite interactions were quantified between plants and caterpillars as well as between caterpillars and parasitoids. Additionally, tri-trophic interactions between plants, caterpillars, and parasitoids were included to capture emergent properties on network structure (Dyer et al. 2010, Pilosof et al. 2017; Figure 3). Alpha diversity of species and interaction at the local-level were quantified within each plot with plot means used in statistical analyses. Local beta diversity represents the turnover of species or interactions and was quantified between plots within each fire return interval category. To estimate standard errors around each estimate we performed bootstrapping with the relative proportions of each species/interaction and the total sample size used to construct 100 simulated plots for each burn category following Chao et al. (2008). Regional-level diversity components were estimated within (alpha) and between (beta) each fire return interval category. Diversity estimates are reported using the inverse Simpson diversity index (1/D) and represent independent measures of alpha and beta following Jost (2007). Documented interactions were used to create and visualize trophic networks for all data and for each fire return interval category (Figure S1).

**Figure 3:**
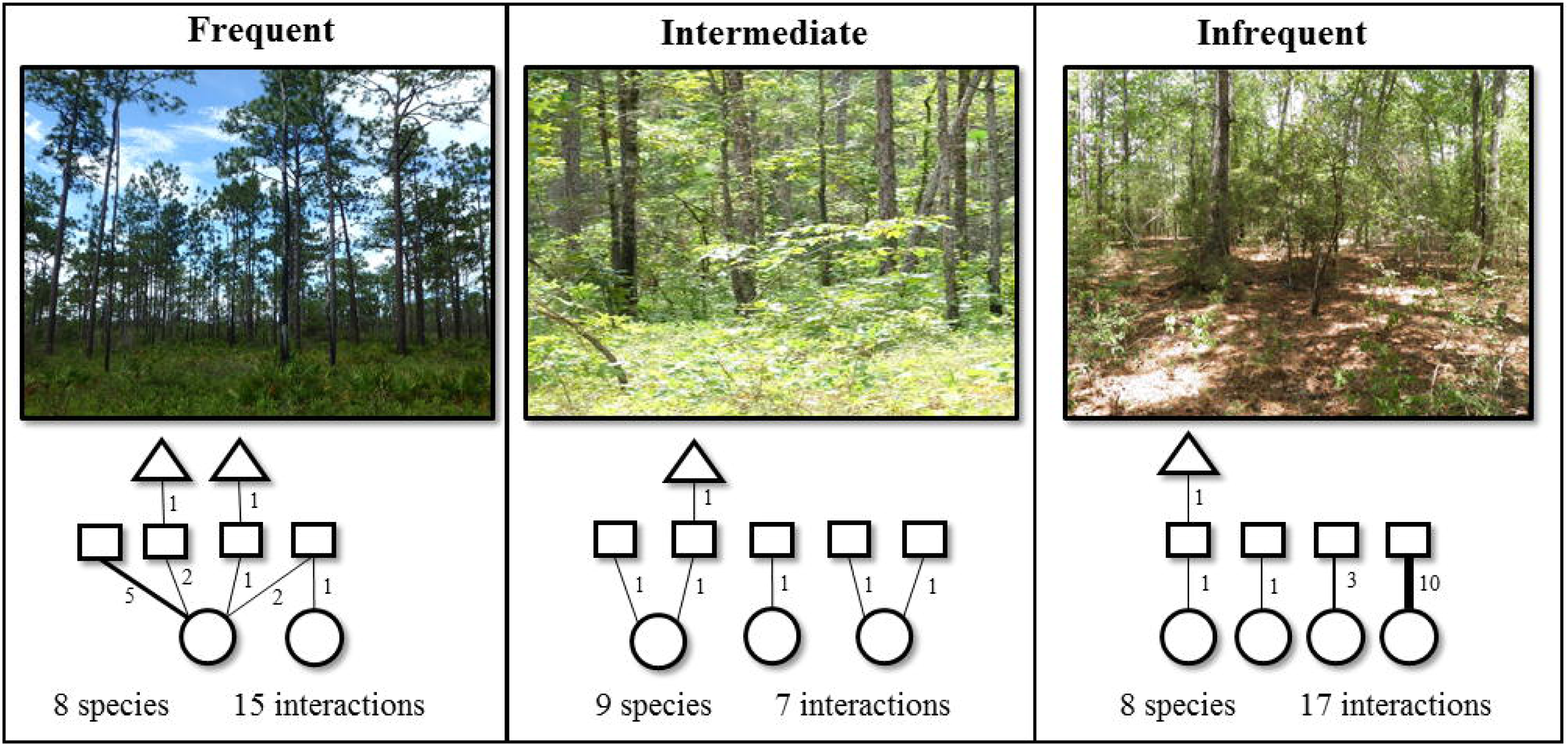
Photo representation of each fire return interval (FRI) category and a sample interaction diversity network found within a single plot in each FRI type. Interactions and their abundances are indicated by solid lines and corresponding numbers, representing interactions between host plants (circles) and herbivores (squares), or between herbivores and parasitoids (triangles), respectively. Each tri-trophic interaction linking a plant, herbivore, and their parasitoid is considered an additional single interaction. Interaction richness and abundance are used to quantify interaction diversity components. Local-scale diversity metrics were calculated with individual plots while regional-scale diversity metrics represent an aggregation of interaction diversity plots and general collections within each FRI category.

### Statistical analyses

To determine if interaction and species diversity across the burn gradient showed similar patterns within plots we utilized analysis of variance, with fire return interval category as an independent variable, and with alpha and beta diversity parameters as response variables. We performed separate univariate ANOVAs for each diversity component at the local scale. Post-hoc analyses utilizing Tukey’s test were performed to identify differences between fire return interval types for each diversity parameter.

To address unequal sampling efforts in terms of number of plots within each fire return interval category, we performed sample-based rarefaction for species and interaction richness. We also calculated nonparametric asymptotic estimators at equal sample coverage levels following Chao et al. (2014) to allow for community comparison across the fire return interval gradient. Discriminant function analyses were conducted to detect differences between species and interactions within fire return interval categories. All analyses were performed in R (v.3.2.3, R Development Core Team 2013).

## Results

The collective sampling effort resulted in a trophic network between 64 host plant species, 183 caterpillar species, and 47 parasitoid species. Combined, there were 1,415 individual interactions between species comprised of 468 unique interactions. 66% of all interactions were detected only once, and only 2% of interactions occurred over 20 times. Within all fire return interval categories, the majority of herbivorous interactions tended to be between one caterpillar and one host plant species. However, some individual plant species were consumed by numerous herbivores and the percentage of caterpillars with more than two host plants had an inverse relationship with time since fire (Table 1). Parasitoid species tended to have a more specialized diet breadth, generally interacting with only a single host species. Each fire return interval category had certain plant species that were involved in a disproportionate number of interactions (Table 1). These highly connected species, such as the host plant *Quercus laevis* (turkey oak) connected to 24% of the entire network, are also referred to as network hubs (Figure S1).

**Table 1:** Relative connectivity of the most linked plant and herbivore species in each tri-trophic network within each fire return interval category. Individual species (i.e. node) connectivity is measured as the percentage of total network links connected to the node in the network. As a summary of diet breadth for each fire return interval category, the percentage of herbivore species that consume 1, 2, or >2 host plants are reported.

| Frequent |  | Intermediate |  | Infrequent |  |
| --- | --- | --- | --- | --- | --- |
| Plant | Connectivity (%) | Plant | Connectivity (%) | Plant | Connectivity (%) |
| *Quercus laevis* | 24 | *Quercus laevis* | 12 | *Vaccinium arboreum* | 7 |
| *Diospyros virginiana* | 12 | *Quercus marilandica* | 10 | *Ilex vomitoria* | 6 |
| *Quercus incana* | 8 | *Quercus margaretta* | 9 | *Quercus laevis* | 6 |
| *Smilax auriculata* | 4 | *Ilex vomitoria* | 6 | *Vitis rotundifolia* | 6 |
| *Vaccinium arboreum* | 4 | *Quercus incana* | 5 | *Vaccinium stamineum* | 5 |
| Herbivore | Connectivity (%) | Herbivore | Connectivity (%) | Herbivore | Connectivity (%) |
| *Gelechiidae* 8 | 3 | *Gelechiidae* 3 | 5 | *Hypeagyris esther* | 3 |
| *Hyperstrovio flaviguttata* | 3 | *Gelechiidae* 10 | 4 | *Noctuidae* 5 | 3 |
| *Gelechiidae* 3 | 3 | *Anisota stigma* | 3 | *Geometridae* 22 | 3 |
| *Erebidae* 1 | 2 | *Megalopyge crispata* | 2 | *Thysanopyga intractata* | 2 |
| *Noctuidae* 5 | 1 | *Hyperstrovio flaviguttata* | 2 | *Gelechiidae* 4 | 2 |
| Host plants per species |  | Host plants per species |  | Host plants per species |  |
| 1 | 70% | 1 | 80% | 1 | 72% |
| 2 | 10% | 2 | 11% | 2 | 22% |
| >2 | 20% | >2 | 9% | >2 | 6% |

### Large scale patterns

Dividing the entire network into regions of similar fire return intervals: frequently, intermediately, and infrequently burned yielded variable patterns in both species and interaction diversity (Table 2, Figure 4). At this larger scale, species alpha diversity increased with longer fire return intervals. However, frequently burned areas had the greatest parasitoid and herbivore species diversity. Parasitoids made up 15% of species richness in frequently burned areas while only 8% in the infrequently burned stands (Table 2). The diversity of interactions did not have a clear pattern across the burn gradient with frequent and infrequently burned regions having higher interaction diversity than intermediately burned regions.

**Table 2:** Diversity measures for both species and interactions calculated at the regional and local scales for each fire return interval category. Alpha and beta diversity components were estimated using inverse Simpson’s index (1/D). Variance around local beta values estimated by bootstrapping. Regional-level values are calculated from the aggregation of plot and general collection data within each fire return interval category. Bracketed values in regional species richness represent plants (P), herbivores (H), and parasitoids (E), respectively. Local level means are reported (+/-SE). Letters denote significant differences between local fire return interval categories based on Tukey’s test at P < 0.05.

| Scale | Fire Return Interval | Species Diversity |  |  | Interaction Diversity |  |  |
| --- | --- | --- | --- | --- | --- | --- | --- |
| | | Richness<br>[P/H/E] | $\alpha$ | $\beta$ | Richness | $\alpha$ | $\beta$ |
| Regional | Frequent | 170<br>[37/108/25] | 14.11 |  | 245 | 29.83 |  |
|  | Intermediate | 115<br>[30/67/18] | 12.28 | 2.11 | 143 | 8.93 | 2.80 |
|  | Infrequent | 145<br>[38/95/12] | 28.18 |  | 158 | 20.44 |  |
| Local | Frequent | 6.84<sup>a</sup> (0.59) | 4.39<sup>a</sup> (0.29) | 5.42<sup>b</sup> (0.63) | 5.44<sup>a</sup> (0.54) | 3.97<sup>a</sup> (0.35) | 15.29<sup>b</sup> (1.39) |
|  | Intermediate | 9.33<sup>ab</sup> (1.45) | 7.26<sup>b</sup> (1.20) | 4.27<sup>a</sup> (0.38) | 7.55<sup>ab</sup> (1.39) | 7.13<sup>b</sup> (1.32) | 8.04<sup>a</sup> (0.44) |
|  | Infrequent | 13.57<sup>b</sup> (2.31) | 6.63<sup>ab</sup> (1.37) | 5.57<sup>b</sup> (0.88) | 11.29<sup>b</sup> (2.61) | 5.57<sup>ab</sup> (1.45) | 9.00<sup>a</sup> (0.21) |

**Figure 4:**
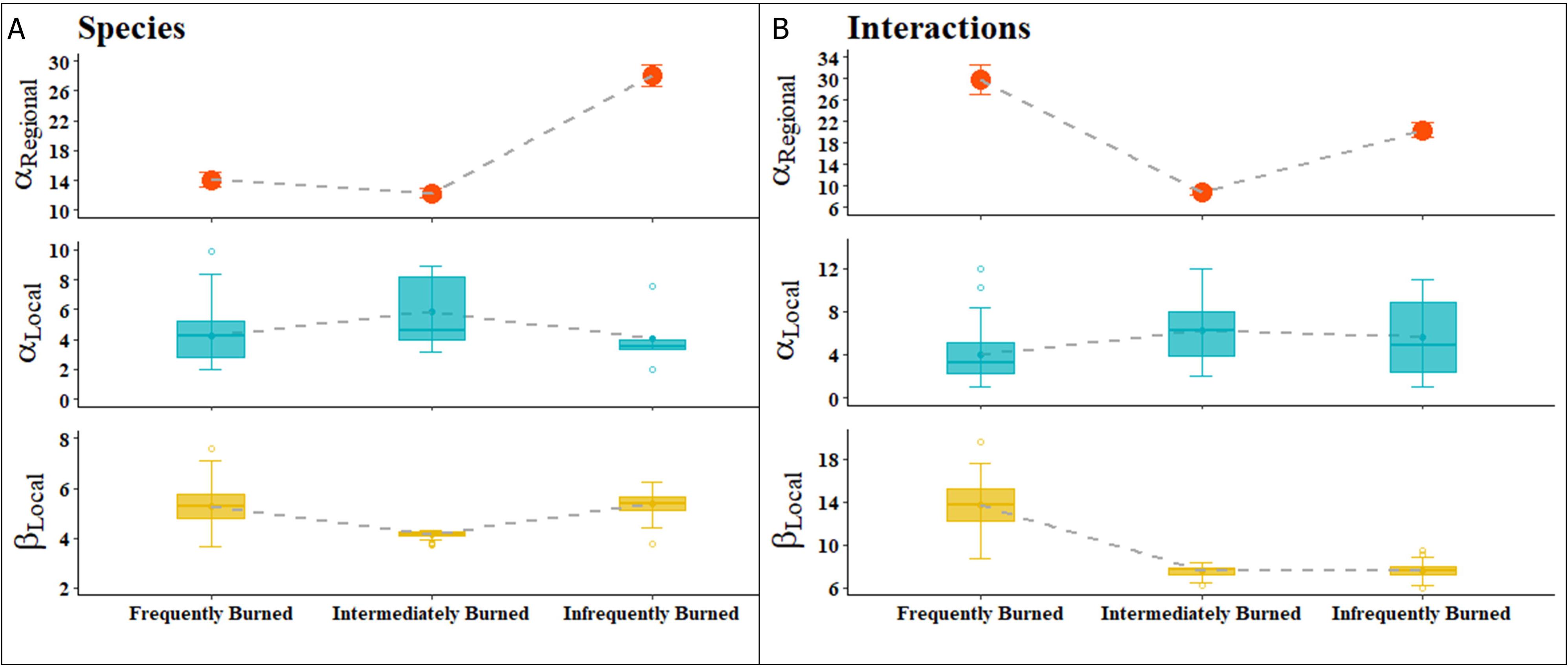
**(A)** Species diversity components and **(B)** interaction diversity components calculated from frequently, intermediately, and infrequently burned sites at hierarchical spatial extents. The broader, regional-level alpha diversity (top panel) represents diversity calculated at each fire return interval category. Our use of the term regional does not infer a singular spatial aggregation, but rather a broader level organization. Local level alpha diversity (middle panel) and beta diversity (bottom panel) represent diversity components calculated at the plot level. Solid lines indicate median value and points connected to dashed lines indicate mean value within local level panels. Diversity values are reported in (1/D).

### Small scale patterns

At the local level, alpha species diversity was significantly higher in intermediately burned plots than in frequently burned plots with infrequently burned plots not significantly different than either (F _(2,63)_ = 6.48, P = 0.003; Figure 4A). Beta species diversity was significantly higher in infrequently burned plots compared to intermediately burned plots but was not different compared to frequently burned plots (F _(2,297)_ = 202.3, P < 0.001). Interaction richness was greatest within infrequently burned plots, while alpha interaction diversity was significantly greater in intermediate burned plots than in frequently burned plots but did not differ between infrequent plots (F _(2,63)_ = 5.03, P = 0.01; Figure 4B). In contrast, beta interaction diversity, or the turnover of interactions, was significantly higher in frequently burned plots, almost double the beta diversity of plots in intermediately and infrequently burned stands (F _(2,297)_= 820.6, P < 0.001).

Rarefaction analyses illustrated that richness of both species and interactions was highest within infrequently burned plots as compared to intermediately and frequently burned plots (Figure S2). Comparing Chao’s asymptotic estimates of species richness at an equal level of coverage of 20 samples, the most species were found within infrequently burned plots (Chao1_infrequent_ = 149) followed by intermediate (Chao1_intermediate_ = 89), and frequently burned plots (Chao1_frequent_ = 78). Interaction richness was also highest in infrequently burned plots compared to intermediately and frequently burned plots in both rarefaction compared at equal sampling effort and comparison of Chao’s asymptotic estimates of interaction richness (Chao1_frequent_ = 75, Chao1_intermediate_ = 110, and Chao1_infrequent_ = 172). Compositional differences between burn interval categories was confirmed through the discriminant function analysis where the first discriminant function explained 99% of the variance and differentiated interactions and species in frequently burned forests from intermediate and infrequently burned forests, with an opposite relationship at the local (species: b = 0.75; interactions: b = 0.83) and regional (species: b = −0.72; interactions: b = −0.64) scale.

## Discussion

We found that the relationship between fire return interval and biodiversity was scale dependent for both species and interactions, as measured by richness, and both alpha and beta diversity components (Table 2). Frequently burned stands were more diverse at a regional-level scale in species and interaction richness as well as interaction alpha diversity. However, these patterns were not consistent when scaling down to the local, plot-level scale. The higher levels of richness among species and interactions, and the higher alpha interaction diversity at local scales in infrequently and intermediately burned stands appeared to be driven by rare species and specialized, single interactions. Shrubby growth forms of hardwood species in longleaf pine forests are maintained by frequent fire, so as fire is removed from the landscape, these species grow and eventually close out the canopy (Hiers et al. 2007). This leads to a depauperate understory of shade tolerant and fire-sensitive plants (Kirkman et al. 2004, Mitchell et al. 2006). As these key plant species are removed due to lack of fire, the increase of fire-sensitive species promotes new interactions.

Frequently burned stands in our study area have more open canopies (Dell et al. 2017), and the characteristic vegetation and associated specialist consumers within more closed canopy stands are not found outside of areas that have not burned in decades as indicated by the segregation of species along the fire return interval gradient. While the assemblages of plants, herbivores, and parasitoids occurring in infrequently burned stands and are characterized by higher richness in comparison to plots that burn more often, interactions between these species are constrained at the local scale. Thus, the lower levels of interaction beta diversity within infrequently burned plots are indicative of the same specialized interactions occurring in individual plots which results in reduced variation in response to disturbance.

One of the most interesting patterns of diversity in the longleaf pine system was the high beta diversity of interactions in frequently burned plots compared to other plots. Lower species richness in frequently burned stands might usually predict similar assemblages in any given plot at the local scale, but this was not the case. While species and interaction richness was lower than in plots without fire, the increased turnover of interactions between plots reveals that stands that burn more often harbor slightly more generalized consumers, an attribute that confers greater potential resiliency to disturbance with increased response diversity (Elmqvist et al. 2003). Fire maintains high response diversity by keeping the ecosystem in a state dominated by longleaf pine and a species-rich, fire-adapted ground cover. In frequently burned forests 20% of the herbivores had a more generalized diet breadth (i.e. > 2 host plant species, Table 1), which provides functional redundancy. In this case, the decreased local alpha diversity can facilitate increased local beta diversity (Chase and Myers 2006), contributing to greater gamma or regional interaction diversity in frequently burned forests - supporting our predictions of frequent fire positively affecting interaction diversity and varying across scale. Focusing in on shared interactions between individual plots illustrates the turnover of interactions between local plots (high β-diversity), this β-diversity summarizes variation in post-disturbance responses regionally, which provides the potential for ecological resiliency.

Furthermore, redundant interactions that may be interchangeable can contribute to sustained ecosystem function (Valiente-Banuet et al. 2015). Higher interaction beta diversity and lower species richness suggest redundant interactions via a rewiring of interacting species in frequently burned forests and may confer resiliency by way of maintenance of ecological function (Lepesqueur et al. 2018). This high degree of interaction turnover may provide an advantage to species adapted to frequently disturbed longleaf pine ecosystem. For example, more generalized diet breadth can be beneficial for individuals post-fire when there is high variability in local plant species composition (García et al. 2016). Response diversity depends on examining multiple spatiotemporal scales to assess full resiliency potential, which may not be evident if only one scale is examined. Regional or ecosystem-level networks represent an aggregation of numerous snapshots in space and time. Thus, there are dynamic processes occurring over time in real networks that are not captured in our static presentation of trophic networks in this system. However, application of a multilayer network perspective allows for associative connections between individual plots (single layer) and the larger scale (multiple layers) by way of shared species and interactions (Pilosof et al. 2017). Therefore, the information we gain by analyzing diversity of interactions are still informative for assessing the impact of fire return interval on the biotic communities. Specifically, contributions to both immediate and long-term resiliencies are found at local and regional-level scales, respectfully.

The relative ecological importance of connectivity in these longleaf networks becomes more apparent when focusing on dynamics of individual species or management of particular species. The relative connectivity of highly connected species, or hubs, has an inverse relationship with fire return interval (Table 1). For example, turkey oak (*Quercus laevis*), a host plant to many herbivores, was represented in 24% of all network links in frequently burned forests compared to only 12% and 6% in intermediate and infrequently burned forests, respectively. While highly connected networks are more resilient to perturbation, the loss of highly connected species would have significant impact on the remaining network and in simulations, eventually leads to network collapse (Bascompte and Jordano 2014). Therefore, maintenance of hub species is an important management consideration. Removal of turkey oak has often been the inappropriate target of intense management in longleaf pine ecosystem (Hiers et al. 2014, Loudermilk et al. 2016). However, our results highlight that the important contributions of turkey oak to functioning networks in longleaf pine forests.

## Conclusion

Disturbances, including natural perturbations such as fire, insect outbreaks, hurricanes, increase habitat heterogeneity which in turn increases the realization of interactions locally and regionally. Variation in interactions is a consequence of varying species abundances, trait distributions and local environmental conditions across the landscape due to variation in disturbance frequency, intensity, duration, and extent. Understanding patterns of interaction diversity within disturbance-dependent networks requires carefully collected data at the appropriate scale at which interactions occur, as well as relevant positions along the disturbance gradient. No biological network is static, and large published webs that are assembled from species inventories (e.g. Bascompte and Jordano 2014) or that examine interactions over large gradients (e.g. Dyer et al. 2007, Forister et al. 2015) are misleading in many ways because identities of interactions often vary across the landscape (Fox and Morrow 1981). At finer scales, such as those examined in our longleaf pine plots, local environmental conditions, community composition, and phenologies differ (Chase and Myers 2006, Garzon-Lopez et al. 2014, Poisot 2015), and the large static network does not exist. Local scale patterns are particularly important in the longleaf pine ecosystem because fine scale heterogeneity in soils, fuels, fire, and dispersal affect plant diversity and community assembly processes (Dell et al. 2017). For example, the processes maintaining assemblages of species and interactions within longleaf pine networks may be deterministic and niche-based at larger scales (entire ecosystems across the landscape) and neutral or stochastic at small scales (1-10m^2^ patches; Dell 2018). Such variable processes across spatial scales, as well as along the disturbance gradient suggest that interactions in this system are governed by both niche and neutral processes as described by the continuum hypothesis (Gravel et al. 2006), presenting an exciting opportunity for future research.

## Acknowledgements

This work was made possible through funding by the Strategic Environmental Research and Development Program of the Department of Defense and the Earthwatch Institute. LD and LR were also supported by DEB-1442103. The funding agencies have no involvement in this study nor publication process.

## Author contributions

LD and LR designed the study; JD, DS, LR, SP, EL, JO and LD collected data; JD, DS and WL analyzed data with LD's input. JD, DS, and LD wrote the manuscript and all authors contributed to revisions.

