## Supplemental Figures for "Scale dependent patterns in interaction diversity maintain resiliency in a frequently disturbed ecosystem"

### 1 Supplementary Figures

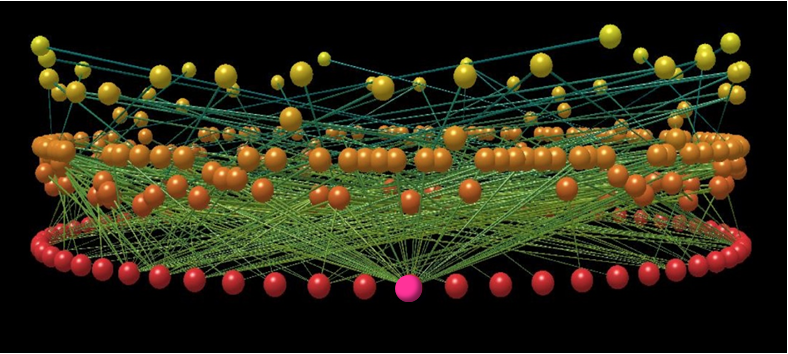
**Figure S1:** 3-D visualization of entire Florida interaction diversity database created using Network 3-D software (Williams 2010, Yoon et al. 2004). The tri-trophic network is comprised of nodes (individual species) and edges (interactions between species). Red nodes represent plants, orange nodes represent herbivores, and yellow nodes represent parasitoid enemies. The pink highlighted plant node represents *Quercus laevis* (turkey oak), a highly-connected network hub.

Williams, R.J. (2010). Network3D Software. Microsoft Research, Cambridge, UK.

Yoon, I., Williams, R.J., Levine, E., Yoon, S., Dunne, J.A., and Martinez, N.D. (2004) Webs on	the Web (WoW): 3D visualization of ecological networks on the WWW for collaborative	research and education. *Vis. Data Anal.*, 5295, 124-132.

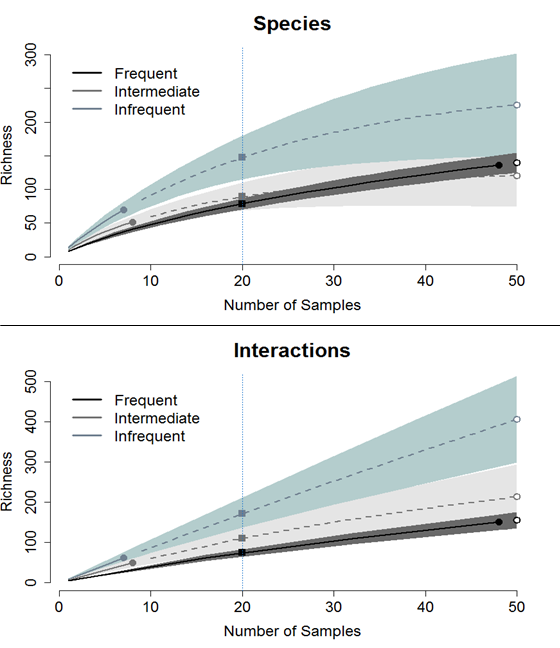

**Figure S2:** Rarefaction curves for species (top panel) and interaction (bottom panel) richness. Solid circles and lines represent observed richness in plots within each fire return interval and open circles represent extrapolated Chao estimates. Solid squares indicate an appropriate level of comparison at the blue dotted line following Chao et al. (2014). Shaded regions represent 95% confidence interval.
